# Cultivar mixtures stabilize wheat baking quality rather than improve it

**DOI:** 10.1101/2025.06.20.660693

**Authors:** Laura Stefan, Silvan Strebel, Karl-Heinz Camp, Sarah Christinat, Dario Fossati, Christian Städeli, Lilia Levy Häner

**Affiliations:** Cultural Techniques and Varieties in Arable Farming, Agroscope, Nyon, Switzerland; Delley Seeds and Plants LTD, Delley, Switzerland; IP-Suisse, Zollikofen, Switzerland; Field-Crop Breeding and Genetic Resources, Agroscope, Nyon, Switzerland; Fresh Food and Beverage Group, Volketswil, Switzerland

**Author notes:** Corresponding author: Laura Stefan.

**Keywords:** baking quality, rheological parameters, wheat cultivar mixtures, quality stability

## Abstract

Integrating diversity into agricultural systems represent a promising way to increase the resilience of crop production. In particular, cultivar mixtures are gaining attention in Europe because they are a practical way to stabilize wheat yields. However, the impact of these mixtures on wheat baking quality remains unclear. In this study, we examined the effects of cultivar mixtures on grain and flour quality. The experiment involved eight Swiss wheat cultivars grown in pure stands, in every possible 2-cultivar mixture, and in the 8-cultivar mixture. The experiment was repeated in eight year-by-site environments, allowing to evaluate the stability of baking quality in mixtures and pure stands.

The results showed that the effects of the mixtures were negative for most flour quality parameters. Furthermore, these effects were not due to changes in cultivar proportions within the mixtures, but rather to cultivar-specific alterations in response to the mixture environment. Finally, mixtures significantly increased the stability of flour quality, by buffering the effects of fluctuating weather conditions.

This study is the first to extensively investigate flour quality in eight contrasting environments. It demonstrates the potential of cultivar mixtures to mitigate the effects of changing abiotic conditions and ensure stable flour quality.

## 1. Introduction

Wheat is one of the most important crops worldwide, contributing about 20% of the total energy intake (Ibba et al., 2022), and providing a variety of staple food products. For wheat baked goods, the quality of the wheat grains and flour is crucial, as it influences the texture and elasticity of the dough, as well as the appearance and taste of the final product (Zanetti et al., 2001). This is particularly important for bread, a product of strong historical, cultural, and religious significance (Das et al., 2023). Key indicators of flour properties, such as protein content, water absorption, and elasticity, are generally influenced by both genetic factors and environmental conditions (Krishnappa et al., 2019). Therefore, fluctuations in environmental conditions, soil types, or disease pressure can lead to significant variations in flour quality across years and locations (Matzen et al., 2019; Torrion & Stougaard, 2017).

In this context, wheat cultivar mixtures are gaining popularity as a potential alternative to the traditional pure stand systems of wheat production (Barot et al., 2017). Indeed, mixtures have been shown to offer several benefits, including improved yield stability (Stefan, Strebel, et al., 2025; Su et al., 2025) or increased resilience to environmental and biotic stresses – such as pests and diseases (Vidal et al., 2017). Therefore, cultivar mixtures could also be a way to improve and stabilize flour quality, by mitigating some of the variability associated with pure stand wheat production. Indeed, the inclusion of several cultivars acts as a buffer against environmental stress, and, in theory, makes it possible to reach a minimum quality threshold thanks to mechanisms of compensation between cultivars (Beaugendre et al., 2024).

However, despite promising potential for wheat quality, most existing research has focused primarily on agronomic traits and yield, leaving a significant gap in our understanding of how cultivar mixtures influence grain and flour quality. The very few studies that have examined the quality of wheat mixtures remain limited and often show inconsistent results (Beaugendre et al., 2024), ranging from positive effects of mixtures on quality (Jackson & Wennig, 1997; Sarandon & Sarandon, 1995) to nonsignificant (Döring et al., 2015; Sammons & Baenziger, 1985) or even negative effects (Cowger & Weisz, 2008). Furthermore, the mechanisms driving such effects are not clear so far: Beaugendre (2024) suggested that two main mechanisms could play a role in driving mixture effects for baking quality, namely proportion shifts – when the harvested proportions of the cultivars in the mixture varies – and component alternation – when plant interactions affect the quality of the component cultivar individually, independently of the final harvested proportions.

In addition, literature has shown that mixture effects on quality are highly dependent on the criteria being assessed. Indeed, quality is a vaguely-defined, multicriteria concept that can include parameters assessed at the grain level (such as grain protein content), flour level (dough elasticity, water absorption potential), but also bread level (bread color, texture, volume) (Fossati et al., 2011). To date, comprehensive assessments of flour quality in cultivar mixtures have been rare. Previous studies often focused on basic quality traits such as protein content (e.g., (Döring et al., 2015; Sarandon & Sarandon, 1995)) or, less frequently, Hagberg’s falling number (e.g., (Cowger & Weisz, 2008)). Only Beaugendre (2024) and Jackson (Jackson & Wennig, 1997) have performed more extensive evaluations, with the former additionally including Zeleny sedimentation value and baking strength (alveograph), and the latter examining bread loaf volume, crumb structure, colour, and texture. Such flour and bread quality analyses are rare, notably because they are expensive in terms of time, personnel and equipment, and they require a large amount of grain/flour to be analysed (at least several kilograms per sample). This can be an obstacle to reproducibility and may explain why there is no previous study evaluating the stability of flour and bread quality in cultivar mixtures across years and/or sites.

Understanding how cultivar mixtures affect grain and flour quality is particularly relevant for Switzerland, where wheat varieties are classified into official quality classes based on baking properties (Fossati et al., 2011). These classes directly influence market prices, with higher quality classes receiving higher prices (swissgranum, 2025). For farmers growing high quality cultivars, it is important to know whether mixing these cultivars will maintain their quality class and, thus, their market value. This uncertainty regarding the quality of mixtures has so far been one of the main barriers to further adoption of cultivar mixtures in Switzerland.

In our study, we aimed to fill this research gap by conducting a comprehensive analysis of grain and flour quality and stability in mixtures and in pure stands in a multi-year, multi-site field trial in Switzerland. We evaluated parameters such as grain protein content, Zeleny sedimentation value, and rheological parameters derived from the extensograph and farinograph analyses. We investigated the effects of cultivar mixtures on grain and flour quality and their impact on Swiss quality classifications. The experimental design further allowed us to assess the stability of grain and flour quality across different environments.

## 2. Materials and Methods

The experimental design used for this study was the same as the one described in (Stefan, Strebel, et al., 2025).

### 2.1 Selected varieties

For the purposes of this study, eight Swiss bread wheat (*Triticum aestivum)* cultivars were selected, which represented all the Swiss bread-making quality classes: Molinera, Bodeli and CH211.14074 with very high quality (called TOP varieties), Schilthorn, Falotta, Campanile and CH111.16373 with high quality (class 1, C1), and Colmetta with medium quality (class 2, C2). All cultivars were winter wheat, except for CH211.14074 which is issued from the spring wheat breeding program, and Campanile which is a summer wheat but issued from the winter wheat breeding program. The cultivars were selected based on their morphological and agronomic characteristics to include differences in yield, protein content, leaf shape and awnness, with no more than 15 cm difference in height and no more than 5 days difference in phenological development to ensure synchrony in maturity. Detailed description of the cultivars is available in Table S1, as well as the allelic profiles of high molecular weight (HMW) and low molecular weight (LMW) glutenin subunits.

### 2.2 Field trials

Field trials were set up over the course of three growing seasons – 2020/2021, 2021/2022 and 2022/2023 – in three sites across the Swiss Central Plateau. The experimental sites were located in Changins (46°19′ N 6°14′ E, 455m a.s.l), Delley (46°55′ N 6°58′ E, 494m a.s.l) and Utzenstorf (47°97′ N 7°33′ E, 483m a.s.l.). Climatic conditions and soil properties for the three sites and three seasons are described in Fig. S1-S3 and Table S2.

Experimental communities consisted of pure stand plots, 2-cultivars mixtures, and one plot with the 8 cultivars mixed. Since 2-cultivar mixtures are most commonly used in Switzerland, they represented the majority of the experimental communities. We sowed every possible combination of 2-cultivar mixtures, amounting to a total of 28 2-cultivar mixtures treatments, to which we added the 8-cultivar mixture. Each community was grown in a plot of 7.1 m^2^ (1.5m*4.7m). We used a complete randomized block design, with 3 replicates, the plots being randomized at each site within each block. Sowing was performed with a small plot drill (Wintersteiger plotseed TC). Density of sowing was 350 viable seeds/m^2^. For the mixtures, seeds were mixed beforehand at a 2x50% ratio for 2-cultivars mixtures and 8x12.5% for the 8-cultivar mixture in proportion to their weight. Plots were sowed mechanically each autumn (see Table S1) and fertilized with ammonium nitrate at a rate of 140 N/ha in 3 applications (40 N/ha at tillering stage/BBCH 22-29; 60 N/ha at the beginning of stem elongation/BBCH 30-31; 40 N/ha at booting stage/BBCH 45-47). The trials were grown according to the Swiss *Extenso* scheme, i.e. without any fungicide, insecticide, and growth regulator.

### 2.3 Data collection

At maturity, we harvested each plot using two different methods: first, we manually harvested a strip of 30 cm × 1.5 m = 0.45 m^2^ along the width of each plot by cutting all individuals (i.e. stems with ears) within the strip right above the ground (see (Stefan, Colbach, et al., 2025) for more details). The harvested strip was located at more than 1 meter from the plot borders to avoid edge effect, and non-homogeneous parts of the plots were avoided. We then air-dried the samples for two weeks. For the mixture samples, we manually sorted the awned stems and ears from the awnless stems and ears. We then separately threshed the awned and awnless individuals and subsequently weighed the grains, in order to obtain the yields from the awned cultivar and from the awnless cultivar. The rest of the plot was then harvested with a combine harvester (Zürn 150, Schontal-Westernhausen, Switzerland).

Protein content (% of dry matter) was measured at the plot level with a near-infrared reflectance spectrometer (ProxiMateTM, Büchi instruments).

Zeleny sedimentation value (ml) was also measured at the plot level based on the ICC standard method 116/1. The analyses were performed by the analytical laboratory of Delley seeds and plants Ltd.

Additionally, flour rheological properties were assessed at the plot level with an extensograph and a farinograph according to ICC standard method 114/1 and 115/1. The analyses were performed by the accredited laboratory “Versuchsanstalt fur Getreideverarbeitung” based in Austria (https://www.vfg.or.at/). The extensograph measures dough resistance to extension (extensograph unit, EU), maximum resistance to extension (extensograph unit, EU), extensibility (mm), and energy (area under curve, cm2). The farinograph assesses the water absorption capacity of the flour (%) and the kneading characteristics of the dough, notably evolution time (min), dough stability (min), and dough softening (farinograph unit, FU). Taken together, we considered 11 quality traits of interest in this study.

### 2.4 Data analyses

All the analyses were performed using R version 4.3.0 (R Core Team, 2019).

Due to missing samples, the grains from the site of Delley in 2023 were not analysed and deleted from the dataset.

#### 2.4.1 Overperformance

In this part, we aimed to evaluate whether the quality of the mixtures was higher or lower than the quality expected from the respective pure stands. For this, we calculated overperformance for each quality parameter as the difference between the observed and expected values of the mixtures, where the expected value is the sum of the values in pure stands weighted by the relative sowing density in the mixture (Stefan, Strebel, et al., 2025):

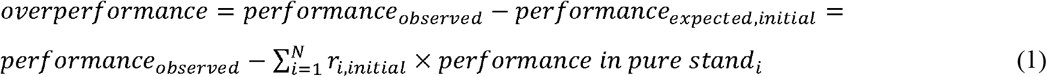

Here, N is the number of cultivars in the plot, and *r_i,initial_*indicates the relative sowing density of cultivar *i* in the mixture. Positive overperformance thus indicates that the mixture had a higher quality value than expected from the pure stands, while negative overperformance suggests that the mixture’s quality was worse than expected from the pure stands. To evaluate whether overperformance was significantly different from zero, we performed t-tests for each quality parameter. These tests were first performed globally, i.e. including all replicates, mixtures, sites and years together for each parameter. Then, we performed additional t-tests (i) per mixture across all sites and years, and (ii) per site and year for all mixtures together. We also tested the effect of class combination (i.e. whether the mixture was TOP-TOP, TOP-C1, etc) on each overperformance parameter using linear mixed models with replication within year and site as random factors, and class combination as fixed effect.

In order to compare changes between parameters, we additionally computed relative overperformance, as

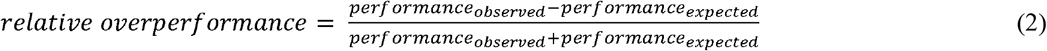

*Proportion shifts*: Finally, to test whether mixture effects were driven by changes in cultivar proportions in the mixture final yield, we calculated the expected performance based on final yield proportions for the mixtures where we could visually distinguish and separate the cultivars:

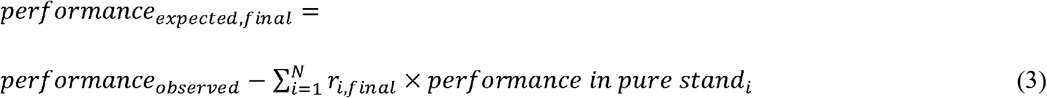

where *r_i,final_*indicates the proportion of harvested grains of cultivar *i* in the mixture. This was only possible for 15 mixtures of 2 cultivars. To assess the role of proportion shifts, we calculated the R^2^ of the expected performance based on final proportions as an estimation of the observed performance, which we compared to the R^2^ of the expected performance based on initial proportions as an estimation of observed performance. Higher R^2^ for expected performance based on final proportions compared to initial proportions would indicate that proportion shifts play a role in driving mixture effects.

#### 2.4.2 PCA and clustering

In order to have a global view of all quality parameters for mixtures and pure stands, we first did a principal component analysis on our 11 quality traits of interest. This was computed using the function *PCA* from the package *FactoMineR (Lê et al., 2008)* on the average parameter values over years and sites. Subsequently, we performed a hierarchical clustering on the results from the PCA, using the function *HCPC* from the same package. The clustering results were visualized using the function *fviz_cluster* from the package *factoextra* (Kassambara & Mundt, 2020).

#### 2.4.3 Stability

Here we aimed to assess the stability of the quality parameters in mixtures and in pure stands. For this, we calculated stability indices for each parameter using WAASB scores, which are the Weighted Average of Absolute Scores from the decomposition of the matrix of BLUPs for genotype x environment interaction effects (Olivoto, Lúcio, da Silva, Marchioro, et al., 2019).

To assess overall quality stability, we computed the Multi-Trait Stability Index (MTSI) developed by Olivoto (Olivoto, Lúcio, da Silva, Sari, et al., 2019), which allows to include several parameters into a single index. The index calculation was based on stability only. The genotype/mixture with the lowest MTSI presents the highest stability for the analysed variables.

The effect of community treatment (i.e. pure stand vs. mixture) on stability indices (i.e. WAASB for each parameter and MTSI) was assessed with linear models. In addition, we also tested the effect of class combination (i.e. whether the mixture was TOP-TOP, TOP-C1, etc) on each stability indices with linear models.

#### 2.4.4. Effects of environmental conditions

The effects of environmental conditions, notably site and year, on grain and flour quality was assessed using linear mixed effects models with year and site as fixed effects, and replication as random factor. This model was run for pure stands and mixtures separately, in order to assess the effects of environmental factors on each crop community. Subsequently, we calculated the proportion of variance explained by either year or site, respectively, by computing partial r-squared.

## 3. Results

### 3.1 Mixture overperformance for grain and flour quality

The results of grain and flour overperformance are summarized in Table 1 and Figure S4. For 10 quality parameters out of 11, the overperformance was significantly different from 0, indicating that the quality of the mixtures was better or worse than expected from the corresponding pure stands. For most parameters, overperformance was negative, which means that the quality of the mixtures was worse than expected from the pure stands. It was only positive for Zeleny sedimentation value and for dough extensibility, indicating a positive mixture effect for these two parameters. Overall, dough resistance, maximum energy, resistance/extensibility and stability had the largest variations between observed and expected values, with up to -16% for the ratio of resistance/extensibility in mixtures.

**Table 1.** Mean overperformance and associated p-value for the grain and flour quality parameters. The overperformance was tested across all sites and years and all mixtures with t-tests. The percentage of change indicates the relative increase or decrease in value from the expected to the observed mixture.

|  | Mean overperformance | p-value | Mean value | % change |
| --- | --- | --- | --- | --- |
| *Grain protein content (%)* | -0.066 | 0.11 | 13 | -0.6 |
| *Zeleny sedimentation value (ml)* | 1.1 | < 0.001 | 52 | 1.7 |
| *Dough resistance (extenso, EU)* | -40.5 | < 0.001 | 297 | -12 |
| *Dough extensibility (extenso, mm)* | 6.23 | < 0.001 | 179 | 3.7 |
| *Dough energy (extenso, cm<sup>2</sup>)* | -6.94 | < 0.001 | 107 | -6.9 |
| *Dough maximum (extenso, EU)* | -47.5 | < 0.001 | 444 | -10 |
| *Dough resistance/extensibility (extenso)* | -0.35 | < 0.001 | 1.7 | -16 |
| *Dough water absorption (farino, %)* | -0.44 | < 0.001 | 60 | -0.7 |
| *Dough stability (farino, min)* | -0.3 | < 0.001 | 1.5 | -12 |
| *Dough softening (farino, FU)* | -5.5 | < 0.001 | 85 | -6.5 |
| *Dough evolution (farino, min)* | -0.17 | < 0.001 | 2.7 | -6.4 |

When looking at the effect of class combination on the overperformance of grain and flour quality, we obtained significant effects only for dough energy and resistance/extensibility ratio (Table S5). Pairwise comparisons with Tukey tests indicated that the mixture effects for dough energy were more negative in C1-TOP and TOP-TOP mixtures compared to C1-C1 mixtures (Figure S5a). Regarding the ratio resistance/extensibility, pairwise comparisons showed that mixture effects were more negative in C1-C2 mixtures compared to C1-TOP and TOP-TOP (Figure S5b).

### 3.2 Proportion shifts

In the mixtures where we could visually distinguish and separate the cultivars, we aimed to determine whether the changes in mixture quality were due to changes in cultivar proportions at harvest. For this we assessed whether we could better estimate performance when accounting for the final proportions of cultivars in the harvested mixture yield. Table 2 shows that the effects of such proportion shifts were dependent on the quality parameter assessed: for instance, proportion shifts did not explain mixture effects in grain protein content, as the R^2^ based on final proportions was equal to the R^2^ based on initial proportions. This was also the case for dough extensibility, resistance/extensibility, water absorption, softening, and evolution. For other parameters, we did observe small effects of proportion shifts, notably for Zeleny sedimentation value, dough resistance, dough energy, maximum, and stability. However, these effects were relatively small, with a maximal increase in R^2^ of 0.03.

**Table 2.** R-squared for estimations of grain and flour quality parameters based on initial sowing proportions and final grain yield proportions. R^2^ is a measure of how much variability within observed performance is explained by the estimations. The higher the R^2^, the better the estimations.

| | $R^2$ of the expected performance based on initial proportions | $R^2$ of the expected performance based on final proportions |
| --- | --- | --- |
| *Grain protein content (%)* | 0.65 | 0.64 |
| *Zeleny sedimentation value (ml)* | 0.65 | 0.67 |
| *Dough resistance (extenso, EU)* | 0.14 | 0.16 |
| *Dough extensibility (extenso, mm)* | 0.51 | 0.50 |
| *Dough energy (extenso, cm<sup>2</sup>)* | 0.26 | 0.29 |
| *Dough maximum (extenso, EU)* | 0.11 | 0.14 |
| *Dough resistance/extensibility (extenso)* | 0.30 | 0.30 |
| *Dough water absorption (farino, %)* | 0.40 | 0.41 |
| *Dough stability (farino, min)* | 0.17 | 0.19 |
| *Dough softening (farino, FU)* | 0.49 | 0.49 |
| *Dough evolution (farino, min)* | 0.46 | 0.46 |

### 3.3 Quality clustering

The principal component analysis followed by hierarchical clustering on all the quality parameters together allowed us to visualise how differently the pure stands and mixtures were behaving. The algorithm partitioned the crop communities into 3 clusters (Figure 1). The grey cluster, on the right side of the graph, is analogous to a TOP quality cluster: it contains the 3 TOP cultivars (211.14074, Bodeli, Molinera) as well as their mixtures (211.14074&Bodeli, 211.14074&Molinera, Bodeli&Molinera). This shows that the mixtures of TOP cultivars remained in the same quality cluster as the pure stands, even though they were closer to the mid-quality cluster in the middle.

**Figure 1:**
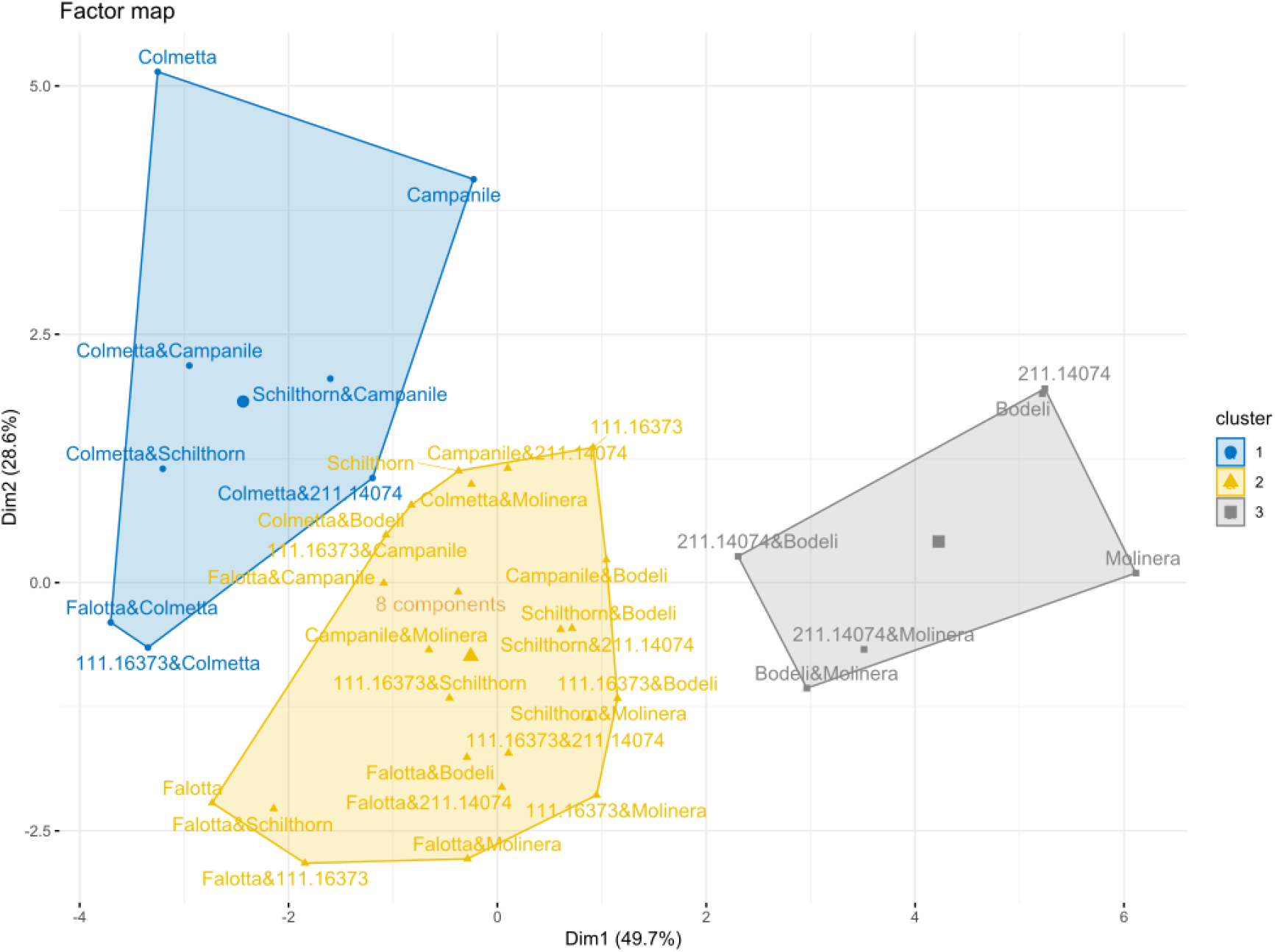
Hierarchical clustering based on principal component analysis of all grain and flour quality parameters. Parameter values were averaged across replicates, years, and sites. Axes indicate the percentage of variation explained. Three clusters were determined by the algorithm: one high-quality cluster in grey on the right, one low-quality cluster in blue on the left, and one mid-quality cluster in yellow in the middle. n =37.

At the opposite side, the blue cluster (on top) could be described as low-quality cluster: it contains Colmetta at its extremity, which is a class 2 cultivar, as well as Campanile, which is a class 1 cultivar. The mixture of these two cultivars can be found in the same cluster, along with 4 other mixtures containing Colmetta and Schilthorn&Campanile. Interestingly, similarly to the high-quality cluster, the pure stands are positioned at the extremity of the cluster, while the mixtures are closer to the center and to the mid-quality cluster. This mid-quality cluster is composed of the pure stands Falotta, 111.16373 and Schilthorn, as well as many mixtures including one of these cultivars. In addition, the mixture with 8 cultivars is also located in the middle of this mid-quality cluster.

### 3.4 Stability of grain and flour quality

The stability of grain and flour quality was assessed across environments (i.e. across sites and years) for each pure stand and mixture. When looking at each quality parameter individually, results showed that mixtures were more stable than pure stands (Figure 2a, Table S6) for all flour quality parameters. For grain quality parameters (that is, protein content and Zeleny sedimentation value), we did not observe any significant effects. When compiling all parameters together into the multi-trait stability index (MTSI), we similarly observed a significant reduction in MTSI score in mixtures compared to pure stands, indicating a significant increase in stability in mixtures compared to pure stands (Figure 2b).

**Figure 2:**
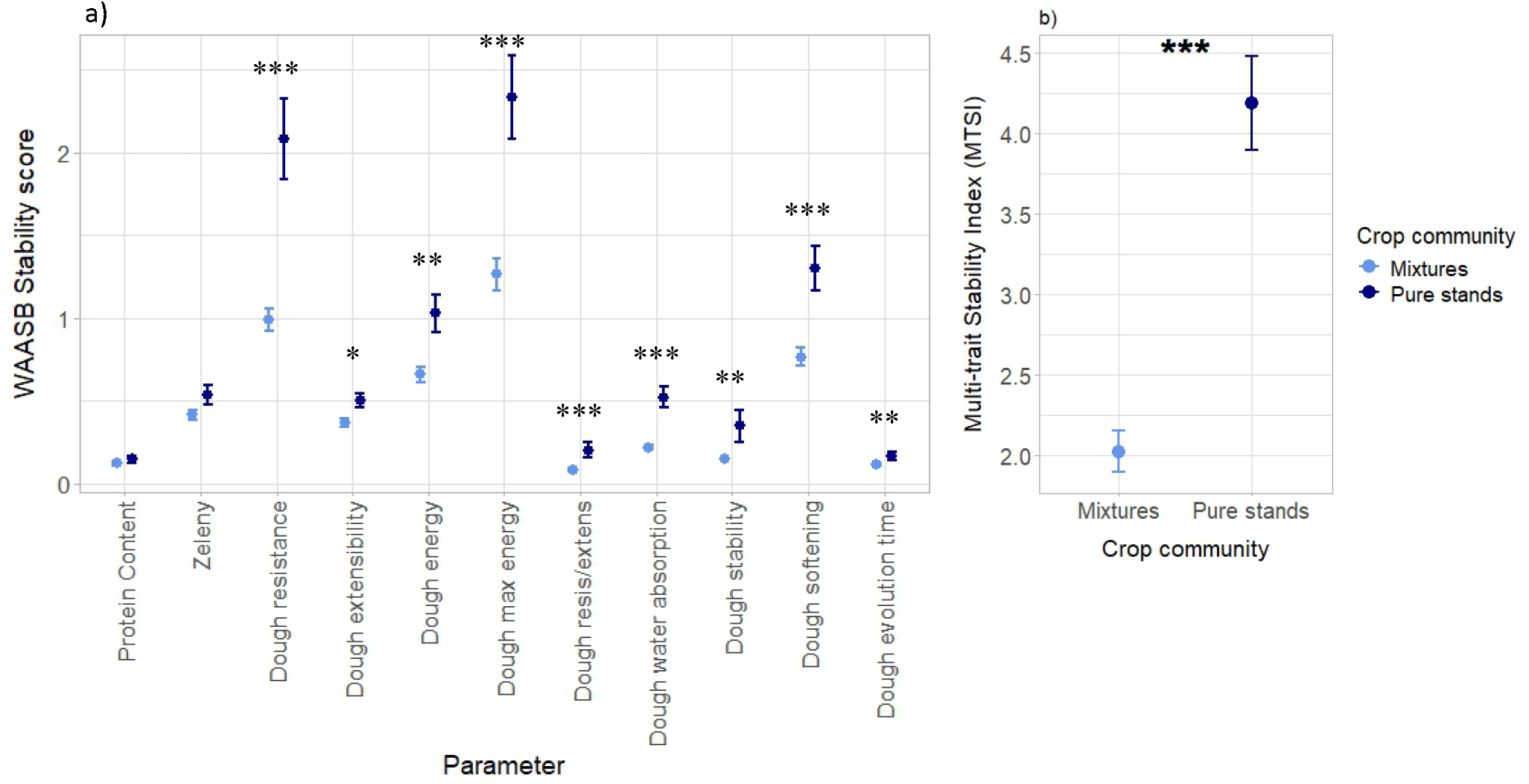
Stability scores for each grain and flour quality parameter (a) and Multi-trait Stability Index (MTSI) in pure stands and mixtures (b). n=37 per parameter. Lower WAASB scores and MTSI indicate higher stability. Dots represent the mean values across plots; lines represent the standard error. Stars placed above or next to the results represent the significance of pure stands vs. mixture. Corresponding statistical results are available in Table S6.

In addition, we looked at the effect of quality class combination on the stability of each quality parameter, as well as on MTSI. We observed significant effects of class combination for several parameters (Table S7), including Zeleny sedimentation value, dough resistance, maximum energy, resistance/extensibility, water absorption, and softening (Figure S7). These results showed that the mixtures combining one TOP cultivar with a C1 cultivar were often the most stable, except for dough softening. Other stable combinations included TOP-TOP, C1-C1, and the 8-cultivar mixture. This was supported by the results for MTSI, for which class combination had a significant effect, and which suggested that the most stable mixtures were C1-C1, C1-TOP, and the 8-cultivar mixture (Figure S8).

### 3.5 Effects of environmental conditions on grain and flour quality

The effect sizes of year and site on the grain and flour quality for pure stands and mixtures are summarized in Table 3. They notably show that environmental conditions could explain up to 37% of the variance, in the case of dough extensibility in mixtures (effect of year). The effect of year was generally larger than the effect of site; this was particularly true for the farinograph parameters. Zeleny sedimentation value was the exception, with a larger effect of site compared to year. For most parameters, we observed a larger effect of environmental conditions in the pure stands compared to the mixtures: this was notably the case for dough resistance, where year and site had a significant effect in the pure stands but not in the mixtures (Figure 3a). This was also observed for dough maximum energy, resistance/extensibility (Figure 3b), water absorption (Figure 3c), softening and evolution. This means that for most quality parameters, pure stands were more sensitive to the effects of environmental conditions than mixtures. Figure 3 in addition shows that pure stands were generally better in absolute terms compared to mixtures, particularly in 2023, where we observed very high values for the flour quality parameters. However, this advantaged decreased in 2021, where the difference between pure stands and mixtures was lower. In the case of water absorption, pure stands even had most of the lowest values in 2021.

**Table 3.** Results of the linear mixed effects models testing the effect of year and site on grain and flour quality, for pure stands and mixtures separately. Effect sizes (F-values) are shown, and stars indicate the significance of the effect. The proportion of variance explained by year/site is also indicated. P-values in boldface type are significant at α = 0.05. * (P < 0.05), ** (P < 0.01), *** (P < 0.001). Green cases indicate where the effect size of year / site is larger in pure stands / mixtures. Orange cases indicate the reverse situation (lower effect sizes).

|  | Pure stands (n = 183) |  |  |  | Mixtures (n = 696) |  |  |  |
| --- | --- | --- | --- | --- | --- | --- | --- | --- |
|  | Year |  | Site |  | Year |  | Site |  |
|  | Effect size (F-value) | Proportion of variance explained | Effect size (F-value) | Proportion of variance explained | Effect size (F-value) | Proportion of variance explained | Effect size (F-value) | Proportion of variance explained |
| *Grain protein content (%)* | 8 ** | 0.16 | 9.5 ** | 0.175 | 8.6 ** | 0.18 | 8.2 ** | 0.18 |
| *Zeleny sedimentation value (ml)* | 1.7 | 0.024 | 6 ** | 0.14 | 1.7 | 0.032 | 9.6 ** | 0.25 |
| *Dough resistance (extenso, EU)* | 12 *** | 0.13 | 11.3 *** | 0.14 | 3 | 0.075 | 2.2 | 0.056 |
| *Dough extensibility (extenso, mm)* | 19.3 *** | 0.2 | 19.4 *** | 0.11 | 37.7 *** | 0.26 | 16 *** | 0.11 |
| *Dough energy (extenso, cm<sup>2</sup>)* | 0.9 | 0.01 | 0.55 | 0.008 | 3.7 * | 0.1 | 1.9 | 0.05 |
| *Dough maximum (extenso, EU)* | 2.1 | 0.02 | 7 ** | 0.09 | 2 | 0.07 | 1.2 | 0.04 |
| *Dough resistance/extensibility (extenso)* | 11.8 *** | 0.16 | 10.3 *** | 0.16 | 8.3 ** | 0.14 | 5.6 * | 0.09 |
| *Dough water absorption (farino, %)* | 30 *** | 0.3 | 3 | 0.03 | 11.8 *** | 0.28 | 2.6 | 0.04 |
| *Dough stability (farino, min)* | 9 ** | 0.117 | 5.5 * | 0.07 | 18.8 *** | 0.245 | 5.8 * | 0.07 |
| *Dough softening (farino, FU)* | 14.4 *** | 0.21 | 6 ** | 0.09 | 15.6 *** | 0.32 | 3.2 | 0.06 |
| *Dough evolution (farino, min)* | 13 *** | 0.18 | 9.8 ** | 0.13 | 12.7 *** | 0.23 | 6.6 ** | 0.11 |

**Figure 3:**
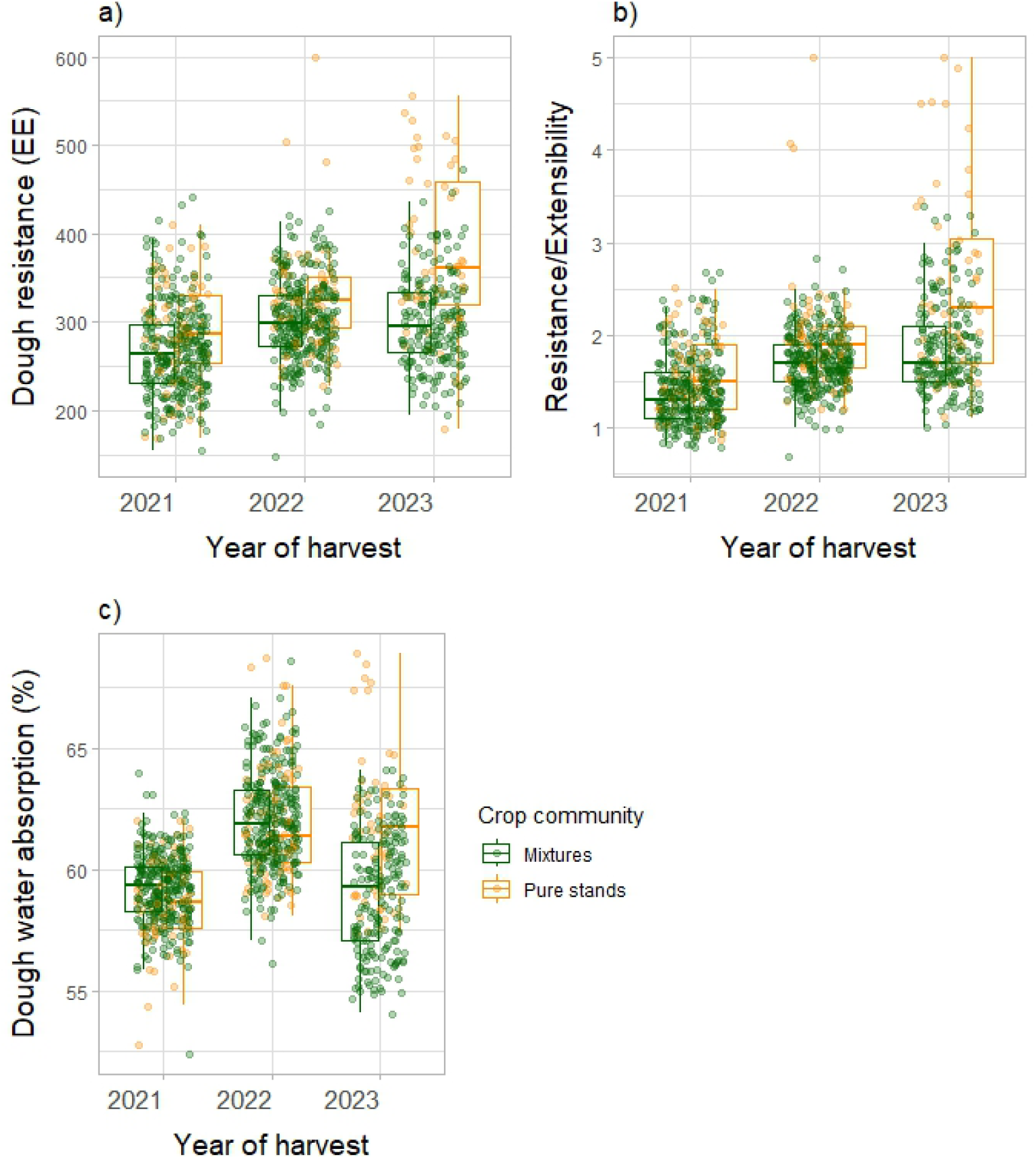
Dough resistance (a), resistance/extensibility (b) and water absorption (c) in response to year of harvest, for pure stands and mixtures. n=950. Horizontal lines represent the median of the data, boxes represent the lower and upper quartiles (25 and 75%), with vertical lines extending from the hinge of the box to the smallest and largest values, no further than 1.5 x the interquartile range. Datapoints are plotted.

## 4. Discussion

In this study, we evaluated the effects of cultivar mixtures on the quality of wheat grain and flour, as well as the stability of this quality across 8 environments. While we had no prior expectations regarding the direction of mixture effects on baking quality, we did expect an increase in stability in mixtures. Grain and flour quality was generally lower in mixtures than in pure stands, and these changes in quality could not be explained by shifts in cultivar proportions at harvest. However, we did observe increased flour stability in mixtures compared to pure stands. Additionally, pure stands were more sensitive to environmental changes than mixtures, suggesting that cultivar mixtures may buffer soil and climatic variations.

### 4.1 Trade-off between quality and stability

Our results showed that, in general, the quality of cultivar mixtures was lower than expected from their respective pure stands, indicating negative mixture effects on quality. This finding aligns with some of the rare previous research on baking quality of mixtures, notably Beaugendre (2024) who found a negative mixture effect on baking strength. However, in contrast to his study, the underlying causes of these quality losses in our mixtures were unclear. Indeed, the contribution of proportion shifts to mixture effects was minor (Table 2), even though changes in cultivar proportions were observed in the harvested grains. Furthermore, proportion shifts cannot explain why mixing two TOP quality class cultivars with similar quality values in pure stands would lead to lower baking quality in mixtures. Rather, we suggest that changes in plant-plant interactions, notably competition between cultivars, can lead to cultivar-specific alterations, such as changes in resource allocation, water uptake, or nitrogen accumulation that can directly impact grain quality (Beaugendre et al., 2024). For instance, in Stefan, Colbach et al. (2025) we showed that cultivar-specific protein content changed in mixtures compared to pure stands. Mixtures have also been shown to alter canopy cover, light interception (Stefan, Strebel, et al., 2025), and microclimate, which can change evapotranspiration, relative humidity, and water loss (Vidal et al., 2017). These alterations can further impact grain quality and grain filling of individual cultivars in mixtures (Torrion & Stougaard, 2017). Investigating such cultivar-specific changes deeper would require measuring the baking quality of each cultivar harvested separately in mixtures, which represents a colossal undertaking, especially since baking analyses require a large quantity of grains.

Lower quality in mixtures was accompanied by an increase in the stability of the quality across the environmental conditions considered in our study (Figure 2), demonstrating a trade-off between quality and stability. This result is not surprising in agronomy, where multiple other trade-offs have already been investigated, such as those between grain quality and grain yield (Anderson et al., 1998) and between productivity and stability (Gutgesell et al., 2024). In our case, we suggest that mixtures may buffer environmental variations through cultivar compensation (Creissen et al., 2013), whereas pure stands are more likely to experience extreme quality values (both high and low) due to their uniform response. Figure 3 illustrates this nicely, showing that pure stands perform extremely well during good years, but also very poorly during wet years, while mixtures tend to perform averagely. Thus, cultivar mixtures reduce the magnitude of GxE effects by averaging cultivar-specific responses to environmental stressors – such as temperature, drought, disease, or heavy rain – and may therefore be particularly well-suited for variable environments, where they would likely perform more predictably than pure stands. This buffering capacity can be viewed as an ecological insurance effect, where diversity within the cropping system provides redundancy and complementary responses to stress (López-Angulo et al., 2023). While this concept has been extensively explored for yield stability in cultivar mixtures (Creissen et al., 2013; Stefan et al., 2024), our findings suggest that it also holds true for quality-related parameters, thereby expanding the benefits of cultivar diversity beyond productivity.

### 4.2 Implications across the wheat value chain

The trade-off that we identified allows to tailor the choice of cultivar or mixture to production goals and environmental conditions. For instance, farmers may choose a pure cultivar for high-end markets in a stable, known environment, or a mixture to ensure lower but more stable quality for broader markets. There are also economic considerations to this choice: is it better to accept lower but stable quality or risk variability for higher potential quality? This question is even more important in the age of climate change, when meteorological variability and the frequency of extreme weather events are expected to increase significantly in the coming decades (Jägermeyr et al., 2021).

Furthermore, specific post-harvest processes could be designed to compensate for the lower average quality of mixtures. For instance, millers already blend cultivars based on the quality of individual components (Cauvain, 2015), so flour from mixtures could easily be incorporated into this process.

Importantly, our results showed that although the quality of the TOP-TOP mixtures was lower, they remained in the TOP quality cluster (Figure 1). Therefore, mixing two TOP-quality cultivars does not result in a degradation of quality class nor in protein content – and, consequently, price – which is an important criterion for farmers (swissgranum, 2025). Overall, our results help shed light on how to classify mixtures based on the quality class of the components. Generally, for mixtures of two cultivars, we found that a mixture’s quality classification is the same as the lowest quality class of its components (e.g., a C1-TOP mixture would be classified as C1; C1-C2 as C2; C2-TOP as C2). These findings could be useful for farmers, millers, breeders, seed suppliers, and institutions responsible for the inscription of cultivars in the official Swiss variety catalogue and the list of recommended varieties. Integrating this classification framework into official guidelines and seed catalogues could indeed help legitimizing the use of mixtures and provide clarity to the wheat value chain stakeholders. Future work should explore the consistency of these results across more genotypes and environments, and assess whether exceptions arise under harsher environmental conditions.

Our study is the first to investigate so many grain and flour quality parameters in eight contrasting year-by-site environments. It clarifies the quality classification of wheat cultivar mixtures and demonstrates their potential to mitigate changing abiotic conditions and ensure a stable flour quality.

## Acknowledgements

We thank Yann Imhoff, Malgorzata Watroba, Reynold Bovy, Julie Roux, and Flavio Foiada for their assistance with field experiments and post-harvest analyses. We also acknowledge the support from Hans Winzeler and Michael Winzeler regarding the choice of the accessions, and the design of the experiment. We are grateful to Versuchsanstalt für Getreideverarbeitung for their support in flour quality analyses and interpretation. This project was jointly funded by the Swiss Federal Office for Agriculture (BLW) and IP-Suisse.

## Funding

This project was jointly funded by the Swiss Federal Office for Agriculture (BLW) and IP-Suisse.

## CRediT authorship contribution statement

**Laura Stefan:** Investigation, Data Curation, Formal analysis, Project administration, Visualization, Writing – Original Draft, Writing – Review and Editing. **Silvan Strebel:** Investigation, Writing – Review and Editing. **Karl-Heinz Camp:** Conceptualization, Funding acquisition, Resources, Investigation, Writing – Review and Editing. **Sarah Christinat:** Conceptualization, Funding acquisition, Writing – Review and Editing. **Dario Fossati:** Conceptualization, Funding acquisition, Resources, Writing – Review and Editing. **Christian Städeli:** Conceptualization, Funding acquisition, Writing – Review and Editing. **Lilia Levy Häner:** Conceptualization, Funding acquisition, Resources, Investigation, Supervision, Writing – Review and Editing.

## Declaration of interests

The authors declare the following financial interests/personal relationships which may be considered as potential competing interests: financial support was in part provided by IP-Suisse and the Swiss Federal Office for Agriculture (BLW). However, the funding partners played no role in the data analyses, generation of the results, and interpretation of the findings.

## Appendix A. Supporting information

Supplementary data associated with this article can be found online.

## Data availability statement

The data will be available on Zenodo upon publication of the manuscript.

## Notes

### Competing Interest Statement

The authors have declared no competing interest.

