## Supplementary Material for "Cultivar mixtures stabilize wheat baking quality rather than improve it"

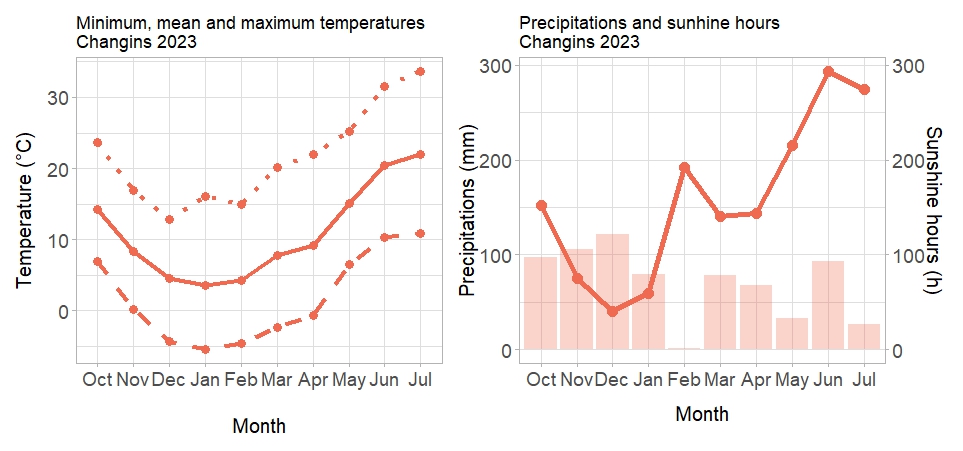

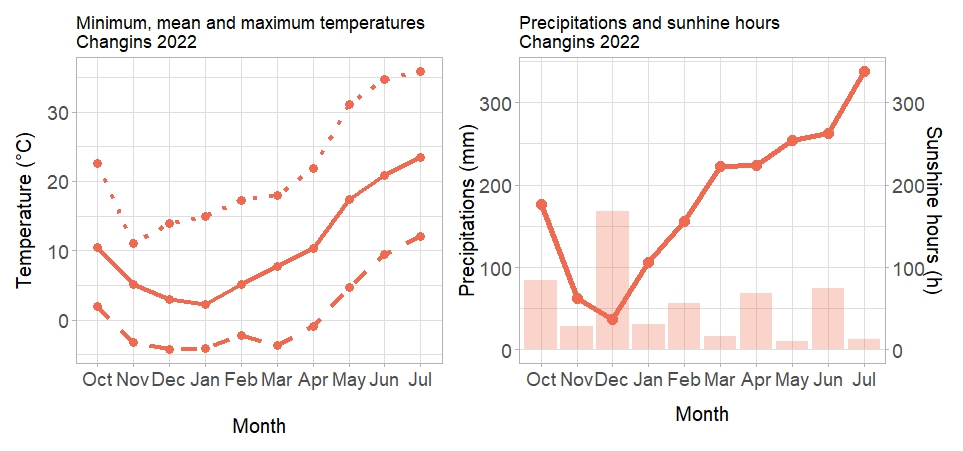

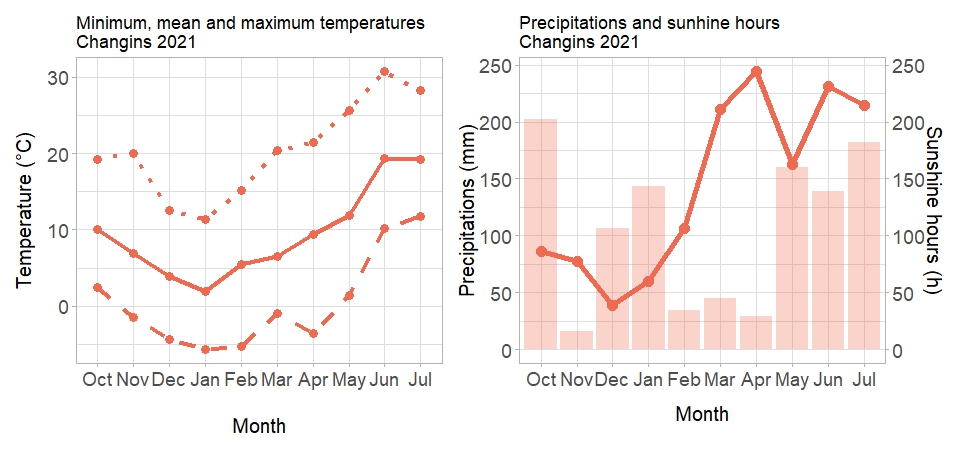

**Fig. S1: Left panel: Minimum, mean, and maximum temperatures in Changins 2021 (a), 2022 (b) and 2023 (c).** Dashed lines represent minimum temperatures, full lines represent the mean, and dotted lines represent maximum temperatures. **Right panel: Precipitations and sunshine hours.** Lines represent the sunshine hours, while bars show the precipitation.

c

**(a)**

c

**(b)**

c

**(c)**

c

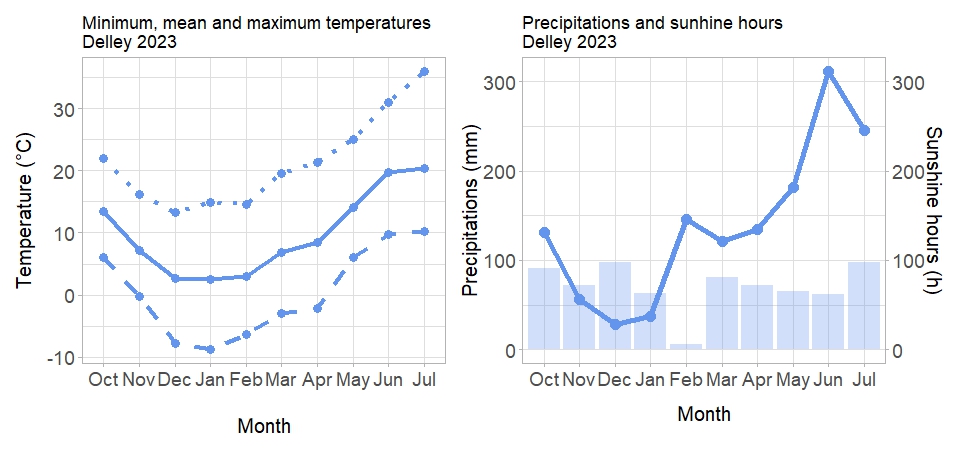

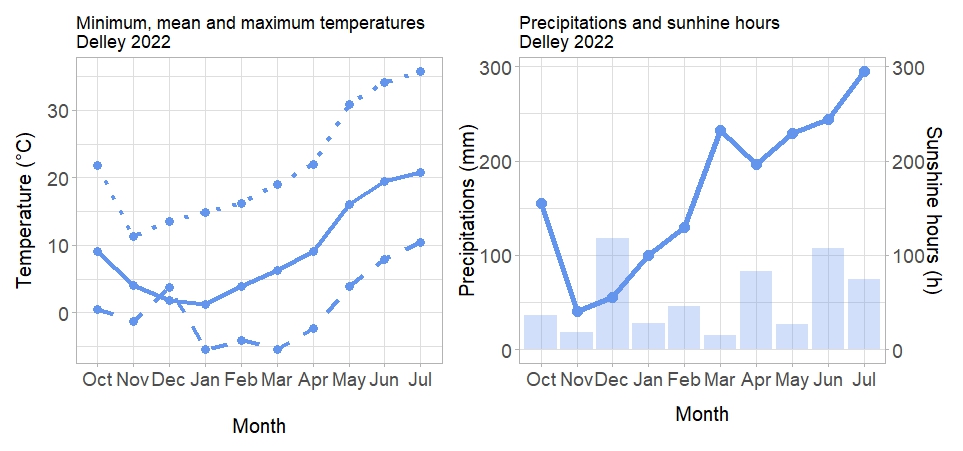

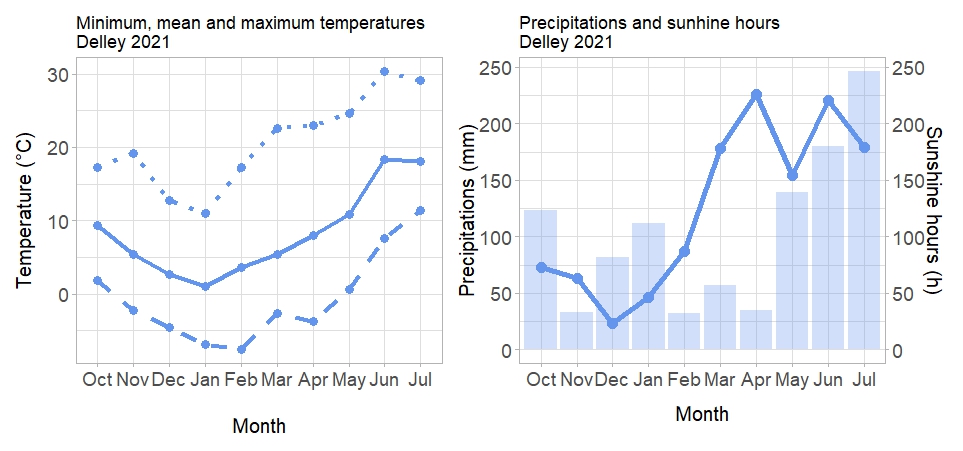

**Fig. S2: Left panel: Minimum, mean, and maximum temperatures in Delley 2021 (a), 2022 (b) and 2023 (c).** Dashed lines represent minimum temperatures, full lines represent the mean, and dotted lines represent maximum temperatures. **Right panel: Precipitations and sunshine hours.** Lines represent the sunshine hours, while bars show the precipitation.

c

**(a)**

c

**(b)**

c

**(c)**

c

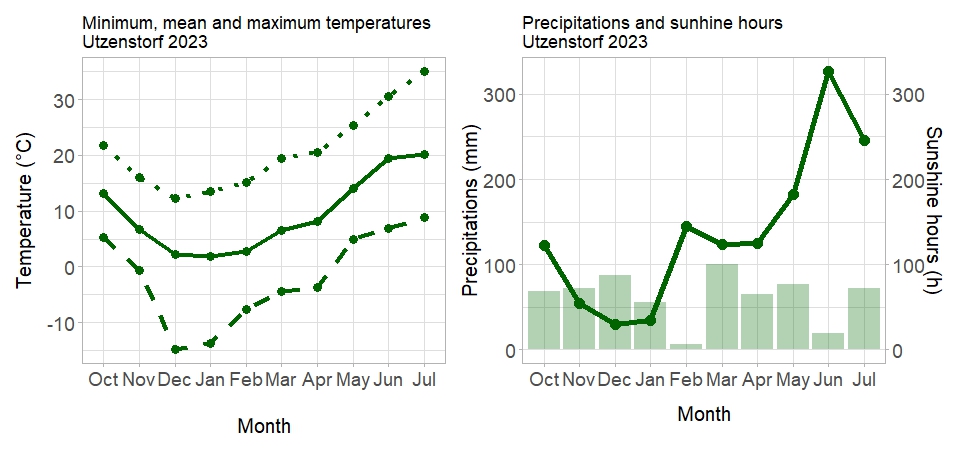

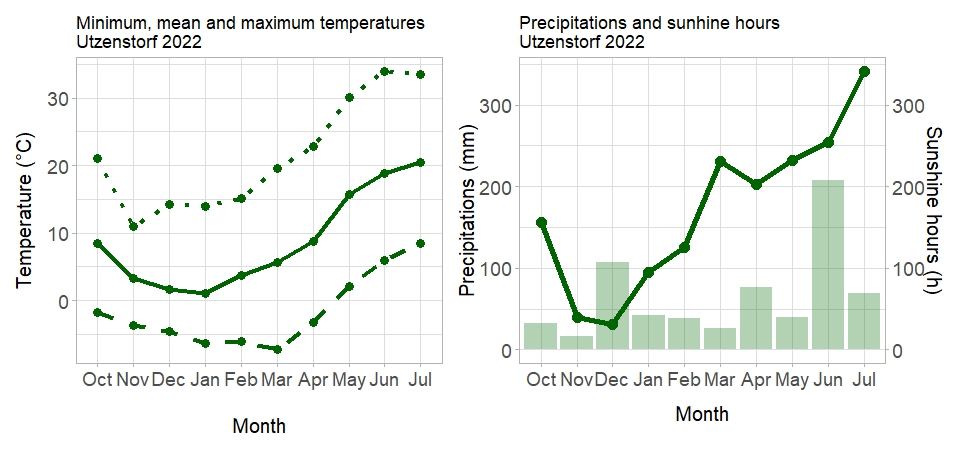

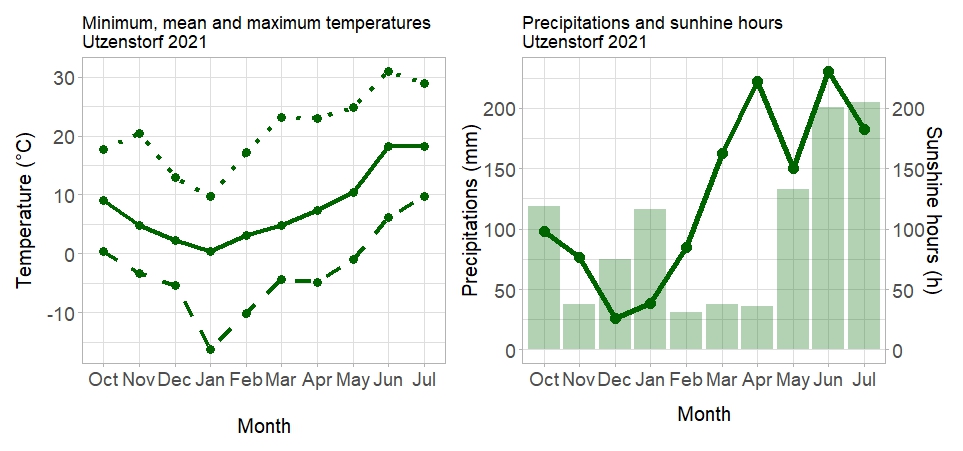

**Fig. S3: Left panel: Minimum, mean, and maximum temperatures in Utzenstorf 2021 (a), 2022 (b) and 2023 (c).** Dashed lines represent minimum temperatures, full lines represent the mean, and dotted lines represent maximum temperatures. **Right panel: Precipitations and sunshine hours.** Lines represent the sunshine hours, while bars show the precipitation.

c

**(a)**

c

**(b)**

c

**(c)**

c

**Table S1: Description of the accessions/varieties used for the experimental trials.**

Data presented here originate from the national variety testing trials and were averaged over years and sites. Relative heading date represents the number of days earlier (in green) or later (in red) than the control varieties. Quality class refers to the national Swiss variety testing system for breadmaking: TOP refers to excellent breadmaking qualities, Class 1 refers to good breadmaking quality, and Class 2 to average. GluA1, GluB1, GluD1: allelic profile for the high molecular weight glutenin subunits. GluA3, GluB3, GluD3: allelic profile for the low molecular weight glutenin subunits.

| Experimental number | Name | Quality Class | Presence of awns | Standardized  Yield (dt/ha) | Zeleny sedimentation rate (ml) | Protein content (%) | Relative heading date | Height (cm) | GluA1 | GluB1 | GluD1 | GluA3 | GluB3 | GluD3 |
| --- | --- | --- | --- | --- | --- | --- | --- | --- | --- | --- | --- | --- | --- | --- |
| 111.15885 | FALOTTA | 1 | Awns | 104.6 | 56.5 | 14.0 | 1.8 | 88.1 | null | 6+8 | 5+10 | a | g | b |
| 111.16373 |  | NA | No awns | 93.2 | 54.3 | 14.4 | 0.4 | 88.2 | 2* | 7+8, 7+9 | 5+10 | a | g | c |
| 111.14470 | COLMETTA | 2 | Awns | 113.0 | 46.5 | 11.9 | -1.4 | 87.5 | 2* | 7+8 | 5+10 | a | c | NA |
| 111.15797 | CAMPANILE | 1 | No awns | 112.0 | 52.8 | 12.9 | 1.2 | 94.4 | null | 14+15 | 5+10 | NA | NA | NA |
| 111.15874 | SCHILTHORN | TOP | No awns | 107.0 | 60.9 | 13.5 | -0.1 | 91.8 | null | 6+8 | 5+10 | a | g | c |
| 211.14074 |  | NA | Awns | 107.0 | 68.5 | 12.7 | -2.4 | 100.7 | 1 | 7+9 | 5+10 | a | g | c |
| 111.15974 | BODELI | TOP | Awns | 100.8 | 62.5 | 14.1 | -0.6 | 96.6 | 1 | 7OE+8 | 5+10 | a | g | c |
| 111.13431 | MOLINERA | TOP | Awns | 89.3 | 62.5 | 15.0 | 0.2 | 87.8 | 1 | 7+8 | 5+10 | a | g | c |

**Table S2: Experimental details and soil status of the fields used for the experimental trials**

|  | *Changins 2021* | *Changins 2022* | *Changins 2023* | *Delley 2021* | *Delley 2022* | *Delley 2023* | *Utzenstorf 2021* | *Utzenstorf 2022* | *Utzenstorf 2023* |
| --- | --- | --- | --- | --- | --- | --- | --- | --- | --- |
| *Sowing date* | 20.10.2020 | 13.10.2021 | 18.10.2022 | 16.10.2020 | 15.10.2021 | 18.10.2022 | 20.10.2020 | 28.10.2021 | 19.10.2022 |
| *Harvest date* | 23.07.2021 | 11.07.2022 | 11.07.2023 | 23.07.2021 | 13.07.2022 | 11.07.2023 | 23.07.2021 | 18.07.2022 | 17.07.2023 |
| *% Clay* | 26 | 22 | 21 | 14 | 20 | 15 | 15 – 20 | 15 – 20 | 15 – 20 |
| *% Silt* | 43 | 47 | 43 | 28 | 46 | NA | NA | NA | NA |
| *% Sand* | 31 | 31 | 36 | 58 | 34 | NA | NA | NA | NA |
| *% Organic Matter* | 2.9 | 2.5 | 2.2 | 1.5 | 2.2 | 1.5 | 2 – 4.9 | 2 – 4.9 | 2 – 4.9 |
| *pH* | 7.7 | 7.6 | 7.1 | 6.9 | 8 | 6.3 | 6.4 | 5.7 | 6.9 |

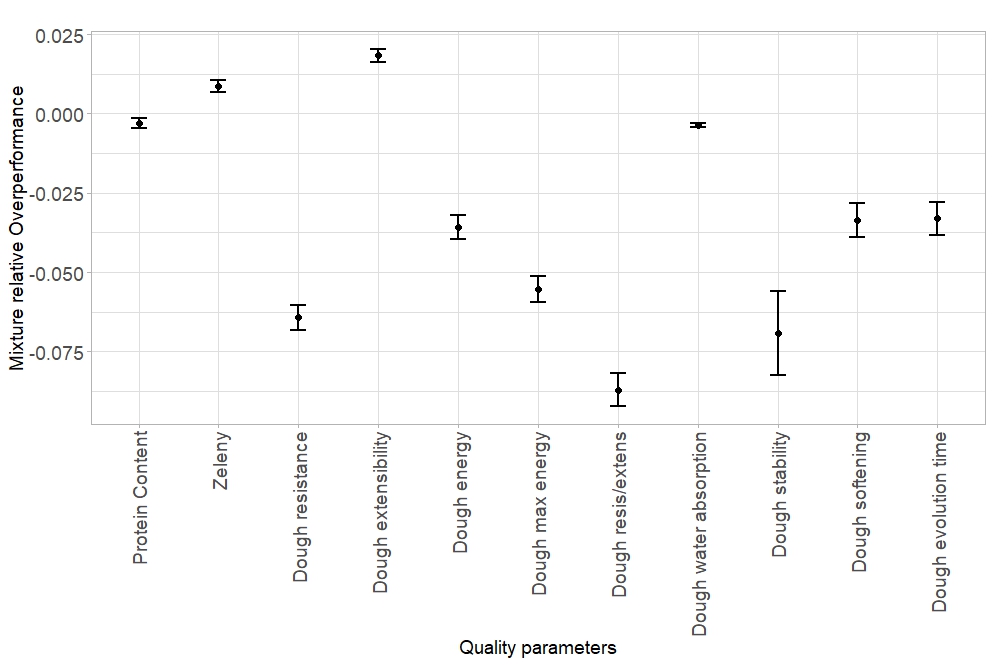

**Figure S4**: Mixture relative overperformance for each of the grain and flour quality parameter assessed. Positive relative overperformance indicates that the mixture was performing better than expected from the corresponding pure stands, while negative relative overperformance indicates that the mixture was performing worse than expected. All results were significantly different from zero, except protein content (Table 1).

Dots represent the mean values across replicates, sites, years, and mixture combinations; lines represent the standard error. n = 673.

|  | Protein content | | Zeleny | | Dough resistance | | Dough extensibility | | Dough energy | | Dough max energy | | Dough resis/extens | |
| --- | --- | --- | --- | --- | --- | --- | --- | --- | --- | --- | --- | --- | --- | --- |
| **Mixture** | **Mean** | **p-value** | **Mean** | **p-value** | **Mean** | **p-value** | **Mean** | **p-value** | **Mean** | **p-value** | **Mean** | **p-value** | **Mean** | **p-value** |
| 111.16373&211.14074 | -0.05 | 0.85 | -0.55 | 0.61 | -59.57 | < 0.001 | 7.63 | 0.08 | -13.59 | 0.02 | -82.08 | < 0.001 | -0.45 | < 0.001 |
| 111.16373&Bodeli | 0.27 | 0.20 | 1.10 | 0.22 | -43.24 | < 0.001 | 7.23 | 0.07 | -5.38 | 0.27 | -48.01 | 0.02 | -0.40 | < 0.001 |
| 111.16373&Campanile | -0.14 | 0.44 | 1.11 | 0.24 | -45.93 | 0.01 | 8.67 | 0.06 | -1.34 | 0.77 | -35.97 | 0.08 | -0.47 | 0.01 |
| 111.16373&Colmetta | 0.23 | 0.48 | 1.29 | 0.43 | -76.14 | < 0.001 | 11.44 | < 0.001 | -9.95 | 0.05 | -80.68 | < 0.001 | -0.70 | < 0.001 |
| 111.16373&Molinera | 0.32 | 0.24 | 0.00 | 1.00 | -48.31 | < 0.001 | 10.25 | 0.01 | -4.64 | 0.33 | -51.49 | 0.02 | -0.44 | 0.01 |
| 111.16373&Schilthorn | -0.06 | 0.77 | 2.08 | 0.03 | -49.10 | 0.01 | 8.19 | 0.08 | -6.18 | 0.21 | -52.16 | 0.02 | -0.43 | 0.02 |
| 211.14074&Bodeli | -0.06 | 0.74 | 1.17 | 0.21 | -46.36 | < 0.001 | 3.81 | 0.30 | -11.69 | 0.06 | -58.98 | 0.02 | -0.32 | 0.01 |
| 211.14074&Molinera | -0.06 | 0.73 | 1.45 | 0.06 | -31.52 | 0.04 | 5.68 | 0.23 | -8.84 | 0.13 | -48.68 | 0.04 | -0.21 | 0.08 |
| Bodeli&Molinera | -0.16 | 0.61 | 0.45 | 0.68 | -52.37 | 0.01 | 4.39 | 0.20 | -14.50 | 0.01 | -75.13 | 0.02 | -0.36 | 0.02 |
| Campanile&211.14074 | -0.21 | 0.30 | -0.96 | 0.32 | -37.67 | 0.01 | 3.30 | 0.40 | -8.24 | 0.16 | -46.80 | 0.05 | -0.31 | 0.02 |
| Campanile&Bodeli | -0.19 | 0.46 | 1.59 | 0.14 | -58.00 | < 0.001 | 7.46 | < 0.001 | -10.33 | 0.01 | -69.35 | < 0.001 | -0.49 | < 0.001 |
| Campanile&Molinera | -0.75 | < 0.001 | 0.89 | 0.30 | -52.36 | < 0.001 | 5.64 | 0.09 | -13.52 | < 0.001 | -71.66 | < 0.001 | -0.40 | < 0.001 |
| Colmetta&211.14074 | -0.14 | 0.53 | 1.09 | 0.40 | -44.41 | < 0.001 | 9.02 | 0.06 | -7.64 | 0.22 | -51.43 | 0.03 | -0.40 | < 0.001 |
| Colmetta&Bodeli | -0.32 | 0.14 | 3.20 | < 0.001 | -45.05 | < 0.001 | 7.70 | 0.03 | -6.98 | 0.14 | -52.14 | 0.02 | -0.43 | < 0.001 |
| Colmetta&Campanile | 0.15 | 0.42 | 1.75 | 0.23 | -40.57 | 0.01 | 6.39 | 0.12 | -5.14 | 0.19 | -43.36 | 0.04 | -0.42 | 0.02 |
| Colmetta&Molinera | -0.23 | 0.15 | 1.77 | 0.04 | -33.34 | 0.01 | 4.57 | 0.11 | -6.75 | 0.13 | -42.20 | 0.04 | -0.30 | < 0.001 |
| Colmetta&Schilthorn | -0.24 | 0.14 | 1.19 | 0.29 | -32.46 | 0.03 | 8.98 | < 0.001 | -0.08 | 0.98 | -26.88 | 0.18 | -0.34 | 0.01 |
| Falotta&111.16373 | 0.24 | 0.26 | 1.60 | 0.06 | -40.21 | 0.03 | 10.61 | 0.03 | -3.43 | 0.39 | -39.49 | 0.09 | -0.41 | 0.02 |
| Falotta&211.14074 | 0.39 | 0.17 | 0.59 | 0.51 | -33.45 | < 0.001 | 6.50 | 0.12 | -8.07 | 0.04 | -45.43 | 0.01 | -0.27 | 0.01 |
| Falotta&Bodeli | -0.01 | 0.96 | 0.65 | 0.47 | -26.94 | 0.00 | 0.35 | 0.93 | -10.71 | 0.02 | -47.15 | < 0.001 | -0.18 | 0.05 |
| Falotta&Campanile | 0.07 | 0.74 | 0.93 | 0.34 | -18.18 | 0.13 | 6.66 | 0.09 | 3.86 | 0.14 | -2.18 | 0.88 | -0.26 | 0.05 |
| Falotta&Colmetta | -0.04 | 0.81 | 1.45 | 0.23 | -33.07 | < 0.001 | 9.73 | < 0.001 | 0.84 | 0.81 | -21.80 | 0.14 | -0.38 | < 0.001 |
| Falotta&Molinera | -0.25 | 0.22 | 1.17 | 0.17 | -28.57 | 0.01 | 7.45 | 0.07 | -5.38 | 0.11 | -38.88 | 0.02 | -0.23 | 0.03 |
| Falotta&Schilthorn | 0.00 | 1.00 | 0.68 | 0.45 | -29.77 | 0.04 | 6.39 | 0.09 | -2.43 | 0.57 | -27.77 | 0.18 | -0.27 | 0.02 |
| Schilthorn&211.14074 | -0.21 | 0.47 | 0.82 | 0.43 | -41.86 | < 0.001 | 5.59 | 0.09 | -8.57 | 0.08 | -47.91 | 0.03 | -0.30 | < 0.001 |
| Schilthorn&Bodeli | -0.40 | 0.08 | 1.32 | 0.22 | -36.66 | 0.02 | 0.91 | 0.72 | -10.66 | 0.02 | -50.82 | 0.04 | -0.25 | 0.03 |
| Schilthorn&Campanile | 0.02 | 0.91 | 0.18 | 0.75 | -22.09 | 0.13 | -2.30 | 0.47 | -6.73 | 0.03 | -29.55 | 0.10 | -0.16 | 0.24 |
| Schilthorn&Molinera | 0.06 | 0.81 | 1.88 | 0.06 | -33.33 | < 0.001 | 2.98 | 0.41 | -9.31 | 0.03 | -50.26 | 0.01 | -0.23 | 0.01 |

Table S3: Mean overperformance and associated p-value for the grain and flour quality parameters. The overperformance was tested across all sites and years for each mixture individually, with t-tests.

|  | Dough water absorption | | Dough stability | | Dough softening | | Dough evolution | |
| --- | --- | --- | --- | --- | --- | --- | --- | --- |
| **Mixture** | **Mean** | **p-value** | **Mean** | **p-value** | **Mean** | **p-value** | **Mean** | **p-value** |
| 111.16373&211.14074 | -0.58 | 0.20 | -0.58 | 0.31 | -11.74 | 0.03 | -0.15 | 0.43 |
| 111.16373&Bodeli | -0.27 | 0.56 | -0.42 | 0.18 | -6.42 | 0.10 | 0.16 | 0.41 |
| 111.16373&Campanile | -0.80 | 0.08 | -0.29 | 0.14 | -3.56 | 0.51 | -0.08 | 0.68 |
| 111.16373&Colmetta | -0.24 | 0.55 | -0.49 | 0.13 | -8.32 | 0.09 | -0.09 | 0.42 |
| 111.16373&Molinera | -0.44 | 0.37 | -0.67 | 0.02 | -14.56 | 0.00 | -0.19 | 0.27 |
| 111.16373&Schilthorn | -0.22 | 0.61 | -0.13 | 0.63 | -2.61 | 0.55 | 0.18 | 0.29 |
| 211.14074&Bodeli | -0.72 | 0.20 | -0.72 | 0.27 | -11.21 | 0.03 | -0.52 | 0.02 |
| 211.14074&Molinera | -0.69 | 0.12 | -0.61 | 0.17 | -2.77 | 0.53 | -0.25 | 0.32 |
| Bodeli&Molinera | -0.87 | 0.22 | -0.22 | 0.41 | -10.16 | 0.07 | -0.47 | 0.04 |
| Campanile&211.14074 | -0.62 | 0.19 | -0.62 | 0.29 | -5.59 | 0.17 | -0.22 | 0.25 |
| Campanile&Bodeli | -1.08 | 0.04 | 0.33 | 0.18 | -4.83 | 0.13 | -0.18 | 0.07 |
| Campanile&Molinera | -1.47 | < 0.001 | -0.18 | 0.59 | -13.05 | 0.02 | -0.47 | 0.06 |
| Colmetta&211.14074 | -0.14 | 0.71 | -0.41 | 0.37 | -6.18 | 0.16 | -0.32 | 0.03 |
| Colmetta&Bodeli | -0.41 | 0.32 | -0.39 | 0.34 | -4.95 | 0.29 | -0.17 | 0.27 |
| Colmetta&Campanile | -0.13 | 0.75 | -0.31 | 0.17 | -2.41 | 0.63 | -0.11 | 0.58 |
| Colmetta&Molinera | -0.58 | 0.27 | -0.17 | 0.53 | -3.23 | 0.33 | -0.31 | 0.08 |
| Colmetta&Schilthorn | 0.04 | 0.92 | -0.31 | 0.09 | -5.33 | 0.13 | -0.01 | 0.94 |
| Falotta&111.16373 | 0.13 | 0.74 | -0.35 | 0.08 | -8.99 | 0.05 | -0.10 | 0.47 |
| Falotta&211.14074 | 0.04 | 0.94 | -0.57 | 0.25 | 0.05 | 0.99 | -0.09 | 0.65 |
| Falotta&Bodeli | -0.26 | 0.61 | -0.08 | 0.75 | -1.54 | 0.65 | -0.24 | 0.10 |
| Falotta&Campanile | -0.25 | 0.46 | 0.21 | 0.39 | 7.00 | 0.09 | -0.21 | 0.24 |
| Falotta&Colmetta | -0.12 | 0.67 | -0.05 | 0.75 | -2.59 | 0.45 | -0.17 | 0.21 |
| Falotta&Molinera | -0.72 | 0.15 | -0.38 | 0.08 | -6.98 | 0.12 | -0.24 | 0.20 |
| Falotta&Schilthorn | -0.21 | 0.62 | -0.06 | 0.69 | -3.93 | 0.38 | -0.04 | 0.80 |
| Schilthorn&211.14074 | -0.15 | 0.70 | -0.71 | 0.08 | -8.20 | 0.07 | -0.10 | 0.55 |
| Schilthorn&Bodeli | -0.41 | 0.42 | -0.10 | 0.78 | -5.00 | 0.24 | -0.18 | 0.28 |
| Schilthorn&Campanile | -0.70 | 0.12 | < 0.001 | 1.00 | -3.11 | 0.40 | 0.01 | 0.94 |
| Schilthorn&Molinera | -0.63 | 0.18 | -0.22 | 0.32 | -5.57 | 0.17 | -0.28 | 0.09 |

|  |  | 2021 | 2021 | 2021 | 2022 | 2022 | 2022 | 2023 | 2023 |
| --- | --- | --- | --- | --- | --- | --- | --- | --- | --- |
|  |  | Changins | Delley | Utzenstorf | Changins | Delley | Utzenstorf | Changins | Utzenstorf |
| Protein Content | Mean | -0.20 | -0.09 | -0.14 | 0.36 | -0.41 | -0.16 | -0.05 | 0.09 |
|  | p-value | 0.07 | 0.49 | 0.09 | 0.02 | < 0.001 | 0.21 | 0.67 | 0.35 |
| Zeleny | Mean | -1.86 | 0.95 | 0.01 | 0.62 | 2.16 | 3.55 | -0.29 | 1.80 |
|  | p-value | 0.00 | 0.09 | 0.98 | 0.13 | < 0.001 | < 0.001 | 0.55 | 0.01 |
| Dough resistance | Mean | -9.95 | -45.77 | -21.47 | -78.47 | -17.99 | 20.96 | -113.63 | -41.88 |
|  | p-value | 0.24 | < 0.001 | < 0.001 | < 0.001 | < 0.001 | < 0.001 | < 0.001 | < 0.001 |
| Dough extensibility | Mean | 3.24 | 6.08 | 4.72 | 9.87 | 11.98 | 0.19 | 9.92 | 3.06 |
|  | p-value | 0.32 | < 0.001 | 0.01 | < 0.001 | < 0.001 | 0.91 | < 0.001 | 0.01 |
| Dough energy | Mean | -0.99 | -15.42 | -5.68 | -17.14 | 6.58 | 10.39 | -17.78 | -11.69 |
|  | p-value | 0.70 | < 0.001 | 0.03 | < 0.001 | < 0.001 | < 0.001 | < 0.001 | < 0.001 |
| Dough max energy | Mean | -12.25 | -76.85 | -32.56 | -119.61 | 3.70 | 48.84 | -119.54 | -53.53 |
|  | p-value | 0.31 | < 0.001 | < 0.001 | < 0.001 | 0.62 | < 0.001 | < 0.001 | < 0.001 |
| Dough resis/extens | Mean | -0.09 | -0.30 | -0.15 | -0.65 | -0.23 | 0.11 | -1.06 | -0.29 |
|  | p-value | 0.20 | < 0.001 | < 0.001 | < 0.001 | < 0.001 | < 0.001 | < 0.001 | < 0.001 |
| Dough water absorption | Mean | 0.61 | 1.12 | 0.52 | 0.82 | 0.21 | -0.24 | -3.40 | -2.41 |
|  | p-value | 0.01 | < 0.001 | < 0.001 | < 0.001 | 0.19 | 0.05 | 0.00 | 0.00 |
| Dough stability | Mean | -0.03 | 0.01 | -0.09 | -0.08 | 0.30 | -0.40 | -0.11 | -1.76 |
|  | p-value | 0.47 | 0.91 | 0.47 | 0.50 | 0.09 | < 0.001 | 0.28 | < 0.001 |
| Dough softening | Mean | 0.13 | -8.70 | -7.43 | -11.46 | 6.37 | -3.02 | -1.81 | -14.29 |
|  | p-value | 0.97 | < 0.001 | 0.01 | < 0.001 | < 0.001 | 0.02 | 0.44 | < 0.001 |
| Dough evolution | Mean | -0.06 | 0.10 | -0.16 | 0.18 | 0.18 | -0.50 | -0.34 | -0.65 |
|  | p-value | 0.32 | 0.03 | 0.03 | 0.07 | 0.11 | < 0.001 | < 0.001 | < 0.001 |

Table S4: Mean overperformance and associated p-value for the grain and flour quality parameters. The overperformance was tested across all mixture combinations, but for each year × site individually, with t-tests.

**Table S5:** Results of the analyses of variance testing the effect of quality class combination on the overperformance of each grain and flour quality parameter. p-values in boldface type are significant at α = 0.05.

Df: degrees of freedom. Sum Sq: sum of squares; Means Sq: mean squares; F-value: variance ratio

| Overperformance in: | Sum Sq | Mean Sq | NumDF | DenDF | F value | p-value |
| --- | --- | --- | --- | --- | --- | --- |
| *Grain protein content (%)* | 5.5 | 1.4 | 4 | 583 | 1.4 | 0.22 |
| *Zeleny sedimentation value (ml)* | 119 | 29 | 4 | 585 | 1.6 | 0.18 |
| *Dough resistance (extenso)* | 7585 | 1896 | 4 | 581 | 0.86 | 0.49 |
| *Dough extensibility (extenso)* | 1821 | 455 | 4 | 582 | 1.8 | 0.12 |
| *Dough energy (extenso)* | 5603 | 1400 | 4 | 583 | 4.7 | **< 0.001** |
| *Dough maximum energy (extenso)* | 46008 | 11501 | 4 | 581 | 2.28 | 0.06 |
| *Dough resistance/extensibility (extenso)* | 2.21 | 0.55 | 4 | 582 | 3.2 | **0.013** |
| *Dough water absorption (farino)* | 12 | 3 | 4 | 582 | 1.7 | 0.14 |
| *Dough stability (farino)* | 4 | 1 | 4 | 584 | 0.5 | 0.72 |
| *Dough softening (farino)* | 1650 | 412 | 4 | 585 | 1.2 | 0.31 |
| *Dough evolution (farino)* | 2.5 | 0.64 | 4 | 585 | 1.3 | 0.28 |

**Table S6:** Results of the analyses of variance testing the effect of crop community (pure stands vs. mixtures) on the stability score (WAASB) of each grain and flour quality parameter, as well as on the multi-trait stability index (MTSI). Estimates and p-values are shown. p-values in boldface type are significant at α = 0.05.

**Figure S5**: Mixture overperformance for dough energy (a) and resistance/extensibility ratio (b) for each quality class combination. Positive overperformance indicates that the mixture was performing better than expected from the corresponding pure stands, while negative overperformance indicates that the mixture was performing worse than expected. A more negative overperformance indicates a mixture with a much lower parameter value compared to expected. Significant differences between quality class combinations are indicated with stars.

Dots represent the mean values across replicates, sites, years, and mixture combinations; lines represent the standard error. n = 673.

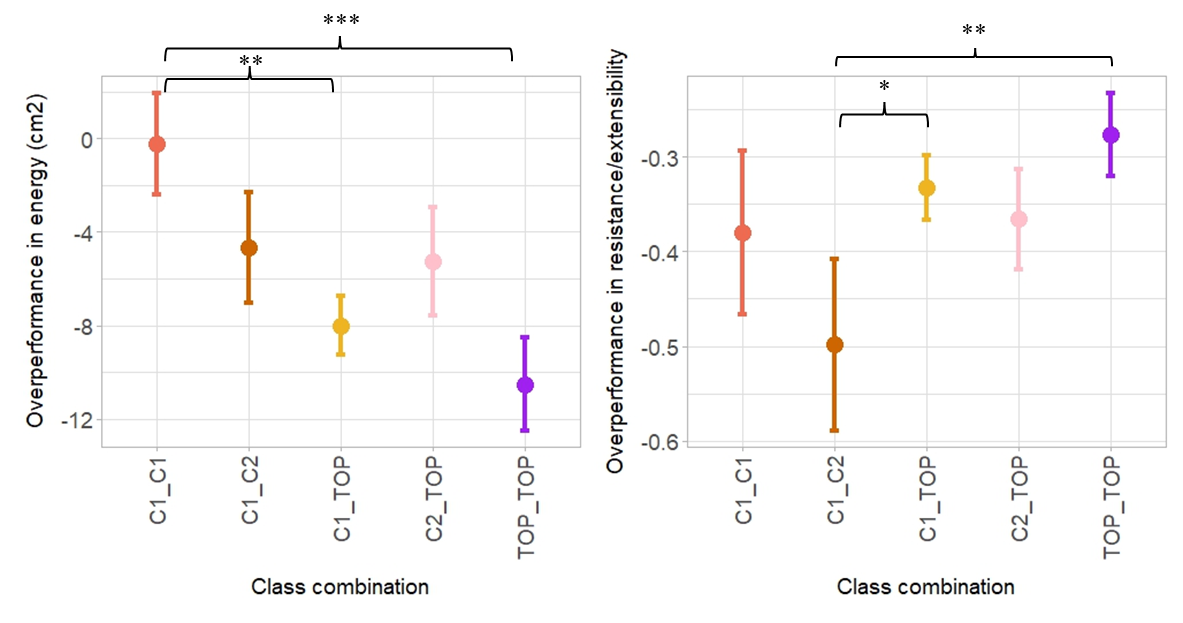

**a)**

**b)**

| WAASB Stability Scores for | Estimate of pure stands vs. mixtures | p-value |
| --- | --- | --- |
| *Grain protein content (%)* | 0.028 | 0.194 |
| *Zeleny sedimentation value (ml)* | 0.12 | 0.062 |
| *Dough resistance (extenso)* | 1.1 | **< 0.001** |
| *Dough extensibility (extenso)* | 0.14 | **0.024** |
| *Dough energy (extenso)* | 0.37 | **0.002** |
| *Dough maximum (extenso)* | 1.07 | **< 0.001** |
| *Dough resistance/extensibility (extenso)* | 0.12 | **< 0.001** |
| *Dough water absorption (farino)* | 0.30 | **< 0.001** |
| *Dough stability (farino)* | 0.20 | **< 0.001** |
| *Dough softening (farino)* | 0.53 | **< 0.001** |
| *Dough evolution (farino)* | 0.05 | **0.006** |
| *MTSI* | 2.16 | **< 0.001** |

**Table S7:** Results of the analyses of variance testing the effect of quality class combination on the stability score (WAASB) of each grain and flour quality parameter, as well as on the multi-trait stability index (MTSI). p-values in boldface type are significant at α = 0.05.

Df: degrees of freedom. Sum Sq: sum of squares; Means Sq: mean squares; F-value: variance ratio

| WAASB Stability Scores for: | Df | Sum Sq | Mean Sq | F-value | p-value |
| --- | --- | --- | --- | --- | --- |
| *Grain protein content (%)* | 8 | 0.02 | 0.00 | 0.62 | 0.75 |
| *Zeleny sedimentation value (ml)* | 8 | 0.44 | 0.06 | 3.10 | **0.01** |
| *Dough resistance (extenso)* | 8 | 8.21 | 1.03 | 4.52 | **0.0013** |
| *Dough extensibility (extenso)* | 8 | 0.22 | 0.03 | 1.22 | 0.33 |
| *Dough energy (extenso)* | 8 | 1.33 | 0.17 | 2.24 | 0.06 |
| *Dough maximum energy (extenso)* | 8 | 10.22 | 1.28 | 4.28 | **0.0018** |
| *Dough resistance/extensibility (extenso)* | 8 | 0.13 | 0.02 | 4.54 | **0.0013** |
| *Dough water absorption (farino)* | 8 | 0.75 | 0.09 | 15.11 | **< 0.001** |
| *Dough stability (farino)* | 8 | 0.31 | 0.04 | 1.93 | 0.09 |
| *Dough softening (farino)* | 8 | 2.70 | 0.34 | 4.08 | **0.003** |
| *Dough evolution (farino)* | 8 | 0.03 | 0.004 | 1.58 | 0.18 |
| *MTSI* | 8 | 33 | 4.1 | 8.1 | **< 0.001** |

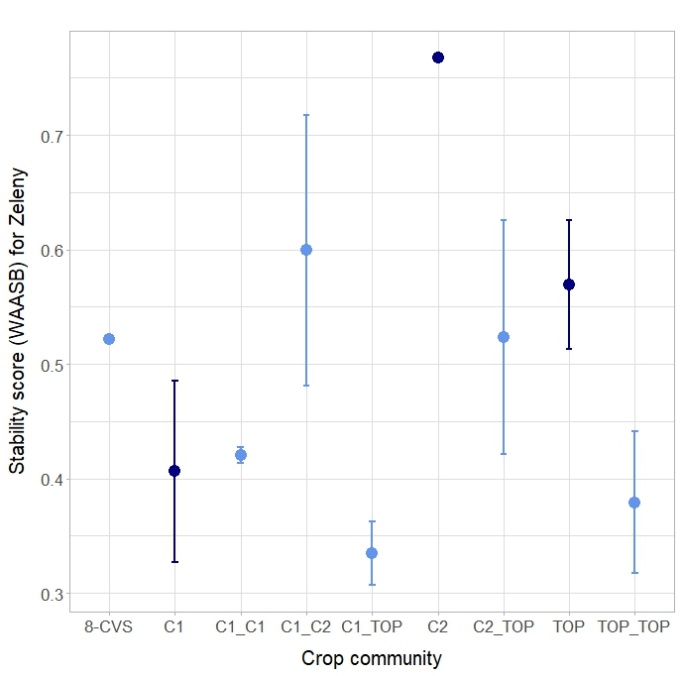

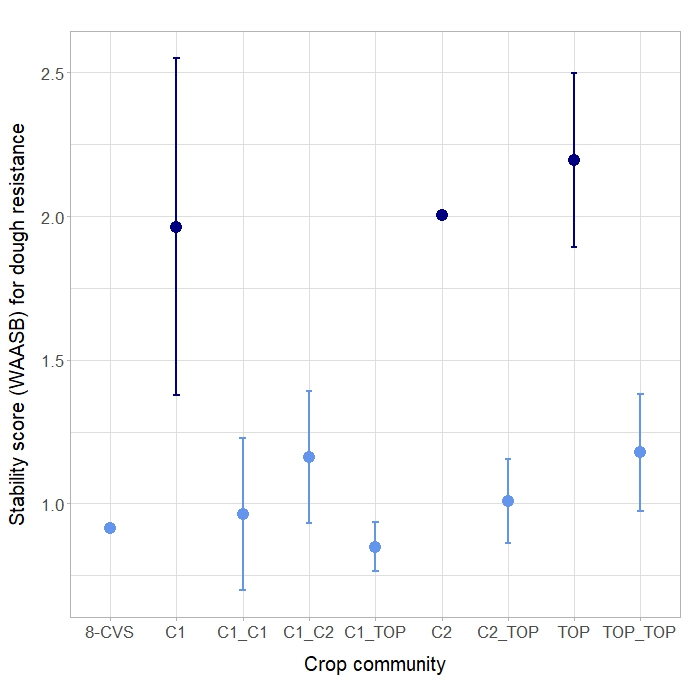

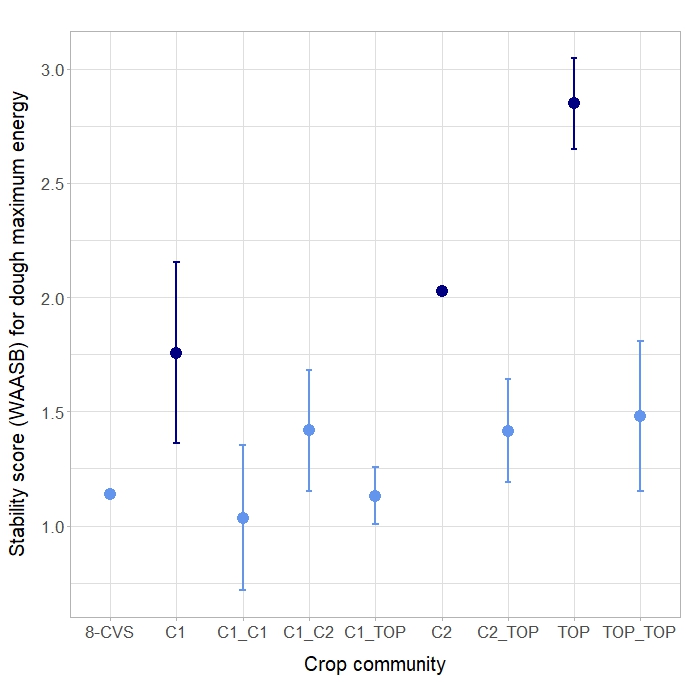

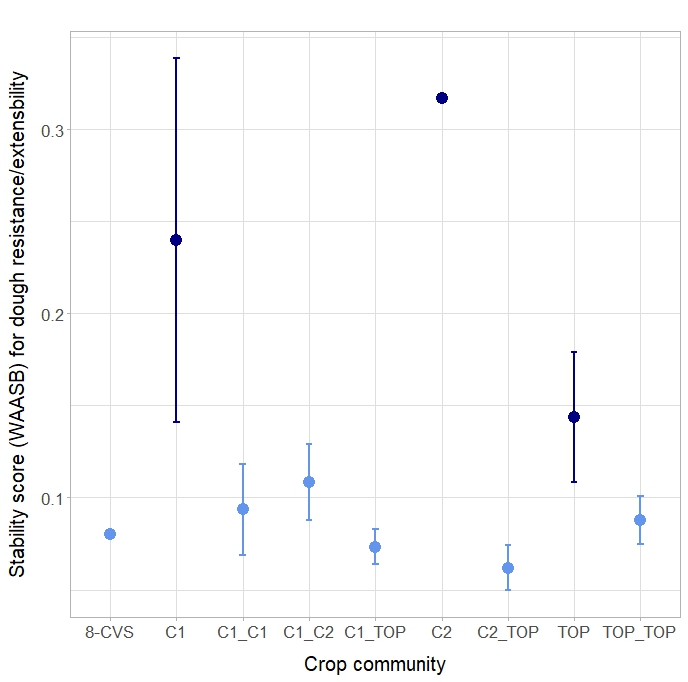

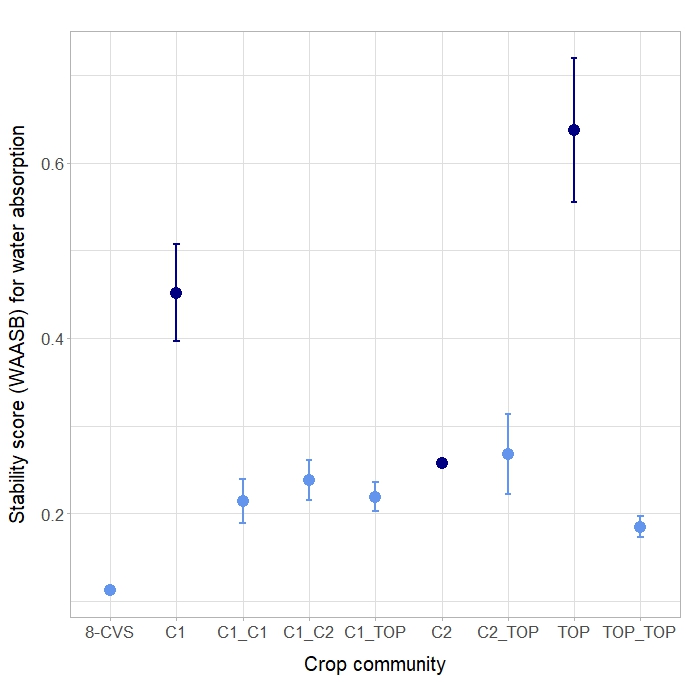

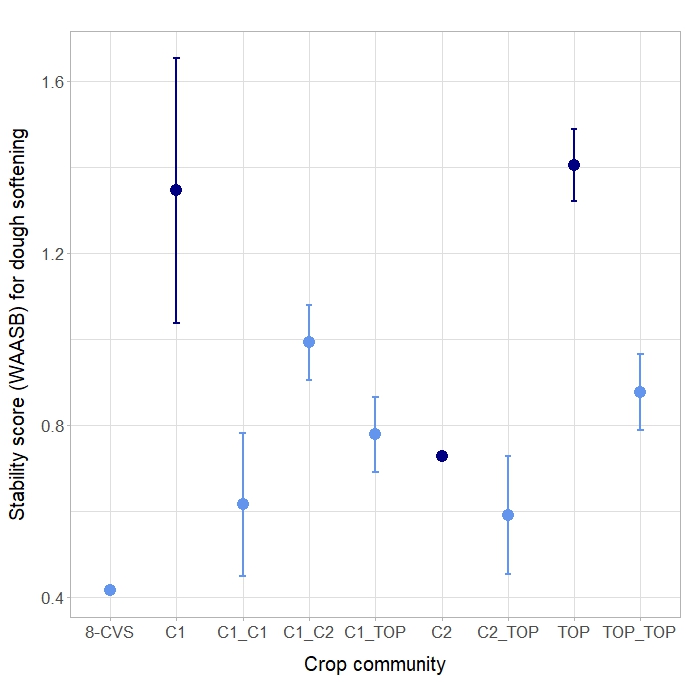

**Fig. S7:** Stability scores for Zeleny (a), dough resistance (b), maximum energy (c), dough resistance/extensibility (d), water absorption (e), and dough softening (f) in response to quality class combination. Dark blue indicates pure stands, while light blue represents mixtures. Dots represent the mean values across plots; lines represent the standard error.

c

**c)**

**d)**

**e)**

**f)**

**a)**

**b)**

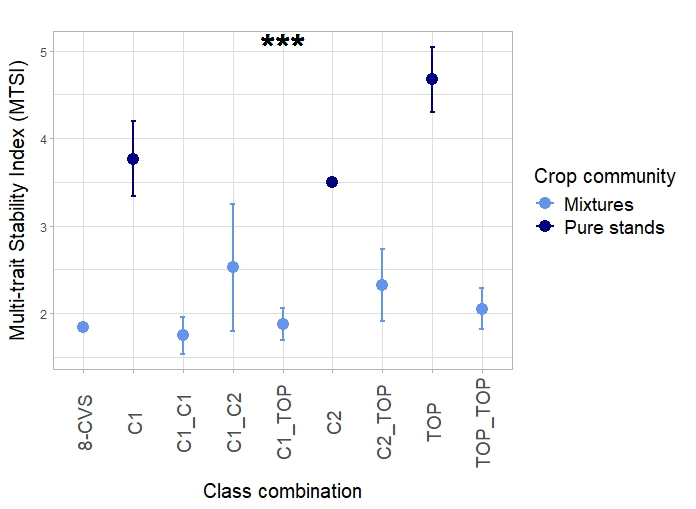

**Figure S8**: Multi-trait Stability Index (MTSI) across class combinations. n=37. Lower MTSI indicates higher stability. Dots represent the mean values across plots; lines represent the standard error. Stars placed above or next to the results represent the significance of class combination. Corresponding statistical results are available in Table S7.
